# Valine Catabolism Drives Bioenergetic and Lipogenic Fuel Plasticity in Prostate Cancer

**DOI:** 10.1101/2024.01.01.573829

**Authors:** Charles L. Bidgood, Lisa K. Philp, Anja Rockstroh, Melanie Lehman, Colleen C. Nelson, Martin C. Sadowski, Jennifer H. Gunter

## Abstract

Metabolic reprogramming is a hallmark of cancer and fundamental for disease progression. The remodelling of oxidative phosphorylation and enhanced lipogenesis are key characteristics of prostate cancer (PCa). Recently, succinate-dependent mitochondrial reprogramming was identified in high-grade prostate tumours with upregulation of enzymes associated with branched-chain amino acid (BCAA) catabolism. We hypothesised that the degradation of BCAAs, particularly valine may play a critical role in anapleurotic refuelling of the mitochondrial succinate pool. Through suppression of valine availability, we report strongly reduced lipid content despite compensatory upregulation of fatty acid uptake, indicating valine is an important lipogenic fuel in PCa. Inhibition of the enzyme 3-hydroxyisobutyryl-CoA hydrolase (HIBCH) also resulted in selective inhibition of cellular proliferation of malignant but not benign prostate cells and impaired succinate production. In combination with a comprehensive multi-omic investigation of patient and cell line data, our work highlights a therapeutic target for selective inhibition of metabolic reprogramming in PCa.

## Introduction

Metastatic castration resistant prostate cancer (mCRPC) is the second leading cause of male cancer mortality, resulting in more than 375,000 deaths globally each year. The objective of current therapies for mCRPC is to inhibit the androgen receptor (AR) signalling axis, which fuels tumour growth and promotes disease progression^1^. Restoration of AR signalling, despite negligible levels of androgens, drives resistance to these therapies, ultimately resulting in treatment failure. The development of resistance is supported by fundamental alterations to energetic pathways in response to the heightened metabolic demand of the tumour, and their metabolic adaptations to therapy^2–4^. Many studies have previously described dysregulated lipid metabolism as a core metabolic adaptation in prostatic malignancy^5–8^. The remodelling of oxidative phosphorylation (OxPhos) has recently been identified as a key characteristic of high-grade prostate cancer tissue. Specifically, these studies describe a metabolic shift from NADH-linked respiration to succinate-linked respiratory function (SLRF)^9,10^. Increased SLRF in high-grade PCa tissue is also coincident with a heightened burden of heteroplasmic mitochondrial DNA (mtDNA) mutations^11^.

Catabolism of the branched-chain amino acids (BCAAs), leucine, isoleucine and valine has been shown to contribute to some pro-oncogenic processes in PCa, but most of what we know about their catabolism comes from their established role as carbon donors during adipogenesis^12–16^. The complete catabolism of the BCAAs produce two major mitochondrial intermediates for tricarboxylic acid (TCA) cycle replenishment: acetyl-CoA and succinyl-CoA. BCAA catabolism is an important fuel source for adipogenesis, demonstrated by significant isotopic carbon labelling (∼40%) of acetyl-CoA from the BCAAs^16^. While the BCAAs contribute to a significant percentage of the cytosolic acetyl-CoA pool in adipocytes, no relationship has yet been established regarding their contribution to the lipogenic phenotype present in PCa. Given the importance of both acetyl-CoA for lipid homeostasis and succinyl-CoA for succinate maintenance, the BCAA catabolic pathway may present an opportune therapeutic target against PCa.

The catabolism of all three BCAAs is universally initiated via the branched-chain amino acid aminotransferases (BCAT1 and BCAT2), but most subsequent downstream reactions are amino acid-specific, allowing interrogation of the role of each individual BCAA. While the BCAAs have been previously identified as a potential target in multiple cancers, their therapeutic potential remains to be investigated in PCa. Here we report that lipid uptake and content of non-malignant prostate cells exhibit a high metabolic dependency on leucine, and that this dependency switches to valine in malignant PCa cell lines. Given the discovery and the importance of succinate metabolism in PCa, we have further investigated the catabolic importance of valine. From the BCAAs, only valine and isoleucine contribute to succinyl-CoA which supplies succinate in the TCA cycle. 3-hydroxyisobutyryl-CoA hydrolase (HIBCH) converts 3-hydroxyisobutyryl-CoA (3HIB-CoA) to 3-hydroxyisobutyric acid (3HIB) and is a central enzyme of valine catabolism. In models of colorectal cancer, specific disruption of valine catabolism via the inhibition of HIBCH directly reduced intracellular succinate levels, inhibited tumour xenograft growth in mice and reduced the occurrence of drug acquired resistance to bevacizumab^17^. In our study, we discovered that BCAAs contribute to the lipogenic phenotype of PCa and demonstrate that metabolic reprogramming of mitochondrial respiration in advanced PCa is associated with increased breakdown of the amino acid valine to fuel the succinate pool and SLRF. Disruption of SLRF via specific inhibition of valine degradation (HIBCH) showed promising therapeutic potential.

## Results

### BCAA uptake and catabolism maintains intracellular lipid supply in prostate cancer

PCa cells are highly lipogenic and *de novo* synthesise fatty acids and cholesterol (lipogenesis). Lipogenesis uses cytoplasmic acetyl-CoA which can be generated from multiple carbon sources, including glucose, fatty acids, acetate and amino acids. BCAAs (leucine, isoleucine and valine) are important carbon fuels for lipogenesis and lipid content in adipocytes. Whether or not BCAAs have a similar role in PCa remains to be elucidated. To evaluate the effects of each BCAA on PCa lipid content, a customised BCAA-depleted medium (BDM) was generated to deprive cells of exogenous BCAAs, to which physiological concentrations of leucine, isoleucine, or valine were re-added. LNCaP, C4-2B and PC3 cells were cultured in BDM for 24 hours before Nile Red staining to quantify intracellular neutral lipid content by quantitative single cell imaging (qSCI) analysis^5^. The absence of all BCAAs resulted in a statistically significant reduction in lipid content within each PCa cell line (***Fig. 1a-c***) indicating that BCAAs potentially contribute to the PCa lipidome. While leucine starvation elicited the greatest reduction to lipids in C4-2B and PC3 cells, valine deprivation caused a comparable reduction in lipid content in androgen-sensitive LNCaP cells. Total BCAA depletion also did not have an additive effect compared to the depletion of any individual BCAA, suggesting the induction of a compensatory mechanism. To confirm whether these effects were from BCAA catabolism, independent from their proteinogenic properties, lipid content was again measured following inhibition of BCAA catabolism by siRNA knockdown of *BCAT1* and *BCAT2 (****Fig. 1f***) in addition to the non-malignant prostatic cell line BPH-1. Significant reductions in lipid content were observed following *BCAT1* knockdown in LNCaP and PC3 cells, while only minor decreases were observed within BPH-1 cells supporting the lipogenic role of BCAAs in PCa cell lines. No significant changes were observed in C4-2B cells (***Fig. 1d, e***). Surprisingly, *BCAT2* knockdown stimulated the accumulation of lipids within all PCa cell lines, while exerting an inhibitory effect on lipids in the benign BPH-1 cell line. This compensatory phenotype was also observable at the transcriptomic level, shown by inverse expression of *BCAT1/2* in response to the suppression of the other (***Fig. 1g***). The strong upregulation of BCAT1 mRNA expression in response to BCAT2 siRNA could have potentially caused an overcompensation in BCAA-fuelled lipogenesis.

**Fig. 1.**
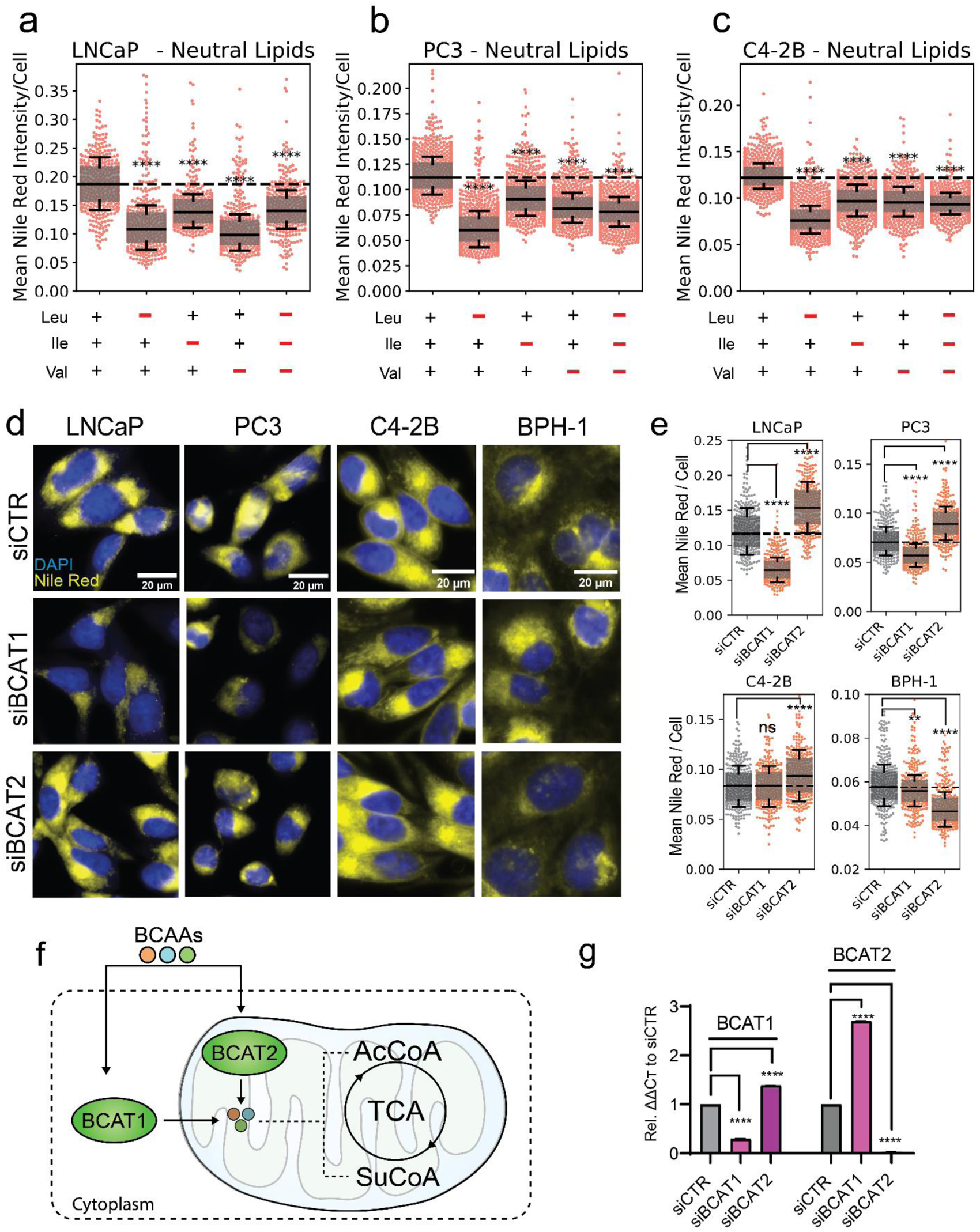
BCAA Uptake and Catabolism is Critical for Intracellular Lipid Maintenance in PCa. (**a-c**) Intracellular neutral lipid content of LNCaP, C4-2B and PC3 cells following 24 hours of exogenous BCAA depletion measured by Nile Red staining and quantitative single cell imaging analysis (qSCI), data representative of 2 independent experiments. (**d**) Representative fluorescent images and (**e**) qSCI analysis of LNCaP, PC3, C4-2B, and BPH-1 cells following Nile Red and DAPI staining after 72 hours of siRNA knockdown of either BCAT1 or BCAT2. Data is representative of 2 independent experiments. (**f**) Schematic of BCAA catabolism by BCAT1 and BCAT2. (**g**) mRNA expression of BCAT1 and BCAT2 following 72 hours siRNA knockdown (siBCAT1 and siBCAT2) measured by qRT-PCR in LNCaP cells. In all experiments, significance was determined by One-Way ANOVA with Dunnett’s Multiple Comparison Test compared to the vehicle control (+BCAAs or siCTR). ns – not significant, **p<0.05, ***p<0.001 ****p<0.0001.

### Exogenous Availability of Valine Co-Regulates Long-Chain Fatty Acid Uptake in PCa

Having established a non-proteogenic, anabolic role for BCAAs in lipid metabolism, exogenous fatty acid uptake was explored as a potential compensatory mechanism in response to BCAA starvation and loss of BCAA-fuelled fatty acid synthesis. PCa cells were cultured in BDM supplemented with and without individual BCAAs at their physiological concentrations. After 24 hours, cells were incubated with the fluorescent palmitate analogue C16:0-BODIPY (C16) (***Fig. 2a***) to measure long-chain fatty acid uptake in response to BCAA depletion. qSCI analysis revealed that both LNCaP and PC3 cells significantly increased C16 uptake in response to BCAA depletion (***Fig. 2d***), while BPH-1 cells significantly reduced C16 uptake. Co-treatment with the non-fluorescent palmitate analogue 2-bromopalmitate (2-BP), a competitive inhibitor of fatty acid transporters (***Fig. 2b***), reduced LNCaP C16 uptake to baseline, and PC3 cell levels below baseline, indicating that C16 uptake was fatty acid transporter-mediated (***Fig. 2d, f***). To probe whether this compensatory response was linked to universal BCAA starvation or was induced by a specific BCAA, C16 uptake was measured in the absence of a single BCAA. Our findings report that valine depletion almost exclusively accounted for the compensatory increase in C16 uptake in malignant LNCaP cells (***Fig. 2c, e***). Notably, benign BPH-1 cells showed this response only after leucine deprivation (***Supplementary Fig. 1***). Together, these findings highlight an important disparity in substrate dependency between benign and malignant prostate cells, most notably a unique response to valine.

**Fig. 2.**
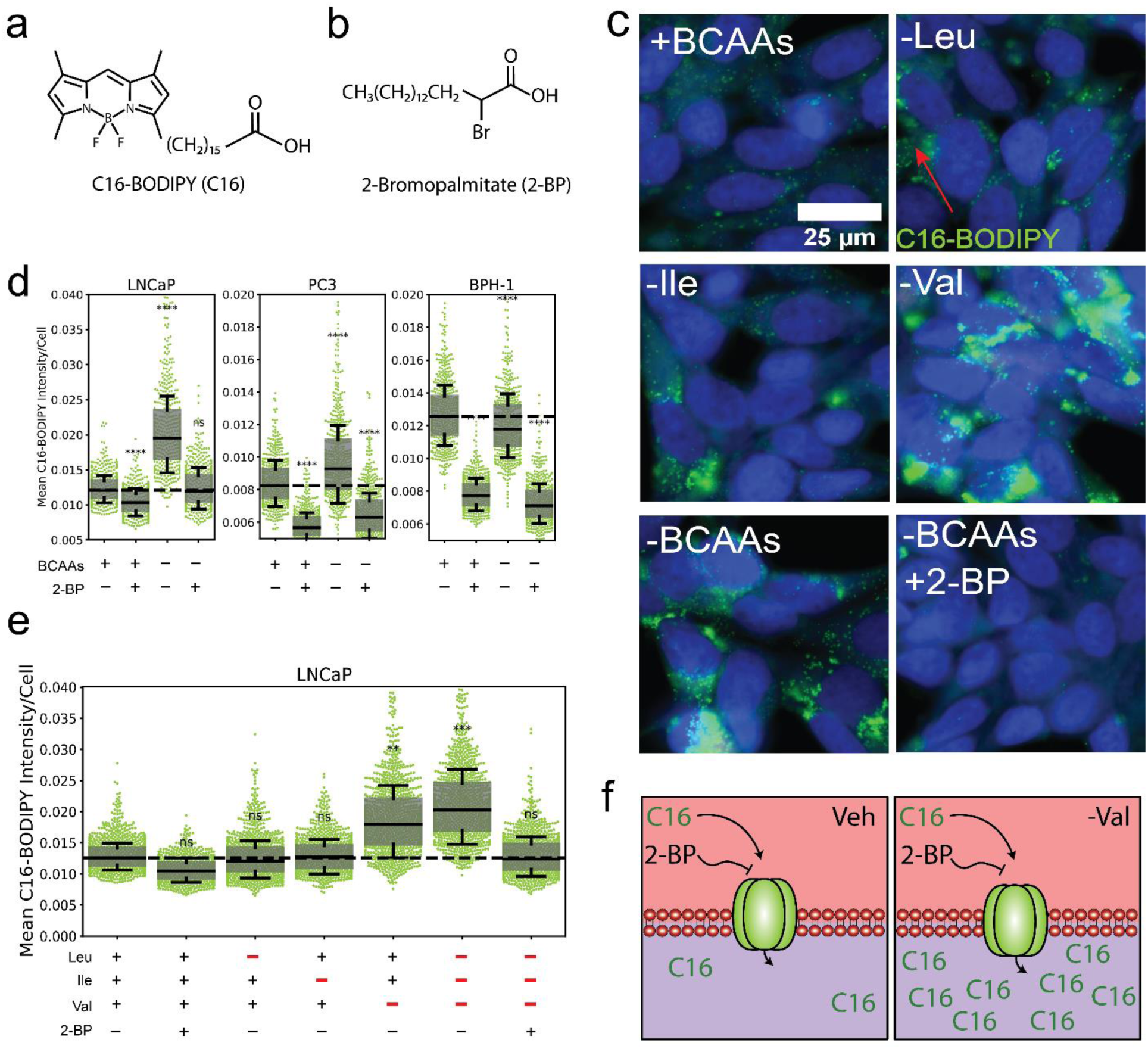
Extracellular Valine Co-Regulates Long-Chain Fatty Acid Uptake in PCa cells. Chemical structure of (**a**) C16-BODIPY (C16) and (**b**) 2-Bromopalmitate (2-BP). (**c**) Representative live-cell fluorescent images and (**d-e**) quantitative single cell imaging (qSCI) analysis of C16-BODIPY (green) uptake following 24 hours of exogenous BCAA deprivation and/or co-treatment with 2-BP in LNCaP cells co-stained with Hoechst 33342. (**f**) Schematic describing valine’s consequence on C16 uptake in PCa cells and its ability to be inhibited by 2-BP. Significance determined by One-Way ANOVA with Dunnett’s Multiple Comparison Test compared to the vehicle control (+BCAAs, -2-BP). ns – not significant, **p<0.01, ***p<0.001, ****p<0.0001.

### Valine catabolism is associated with enhanced succinate-linked respiratory function (SLRF), AR activity and survival across the PCa disease spectrum

To explore the clinical association of valine catabolism and intratumoral SLRF, publicly available patient-derived datasets were interrogated using a customized metabolic gene signature (denoted as MitoS), incorporating genes encoding enzymes necessary to catabolize 3HIB-CoA into succinate, and the respective subunits of SDH (***Fig. 3a***). The MitoS signature was evaluated against an RNA-seq dataset containing matched benign and malignant tissues from 16 localized PCa prostatectomy samples (Schopf et al., 2019 – EGAD00001005931)^11^. The MitoS gene signature was significantly (p=0.029) enriched in malignant versus benign tissue (***Fig. 3b,c***). Finally, the MitoS signature was compared to a previously published gene signature (ARPC) representing AR-high tumours^29^. Linear regression analysis revealed a statistically significant correlation (p=0.0002) between the MitoS and ARPC gene scores within malignant samples, indicating that valine induced SLRF is supported in localised disease and is highly correlated to AR activity (***Fig. 3d***).

To examine the association of valine catabolism in advanced and metastatic disease, we interrogated a microarray (RNA) dataset generated from long-term (21-day) treatment of PCa cells with the AR antagonist, enzalutamide (Tousignant et al., 2020 – GSE143408)^4^. This analysis revealed significantly increased levels of the genes responsible for succinate generation downstream of HIBCH in androgen-sensitive LNCaP cells, suggesting that advanced PCa responds to AR antagonism by further enhancing valine catabolism and SLRF (***Fig. 3e***). To determine the relevance of this mitochondrial phenotype in metastatic castrate-resistant disease, the MitoS signature was compared against a published RNAseq dataset (Labrecque et al, 2019 – GSE126078) of molecularly subtyped patient tumours^29^, which we then stratified for MitoS enrichment. This uncovered a select cohort of AR positive (ARPC) mCRPC patients whose tumours were both highly (MitoS^high^) and moderately (MitoS^med^) enriched for the MitoS gene signature (***Fig. 3f***). Inversely, tumours that possessed neuroendocrine PCa features (NEPC) were negatively associated with these genes, notably HIBCH and succinyl-CoA ligase subunit beta (*SUCLG2*) (***Fig. 3g***). Finally, RNAseq of patient tumours with mCRPC (n=208) were analysed against longitudinal survival data (Abida et al., 2019) to determine whether the MitoS gene signature was predictive of clinical patient outcomes^30^. This analysis revealed that patients with a high MitoS score (top 50%) were associated with a 6.3 month reduced mean survival time (MST) (p<0.0001) than those in the bottom 50%. In concordance with this result, analysis of only *HIBCH* had a 6.8-month reduced MST (p<0.0001) and most notably, the succinate dehydrogenase subunit *SDHB* had a 9.8 month reduced MST.

**Fig. 3.**
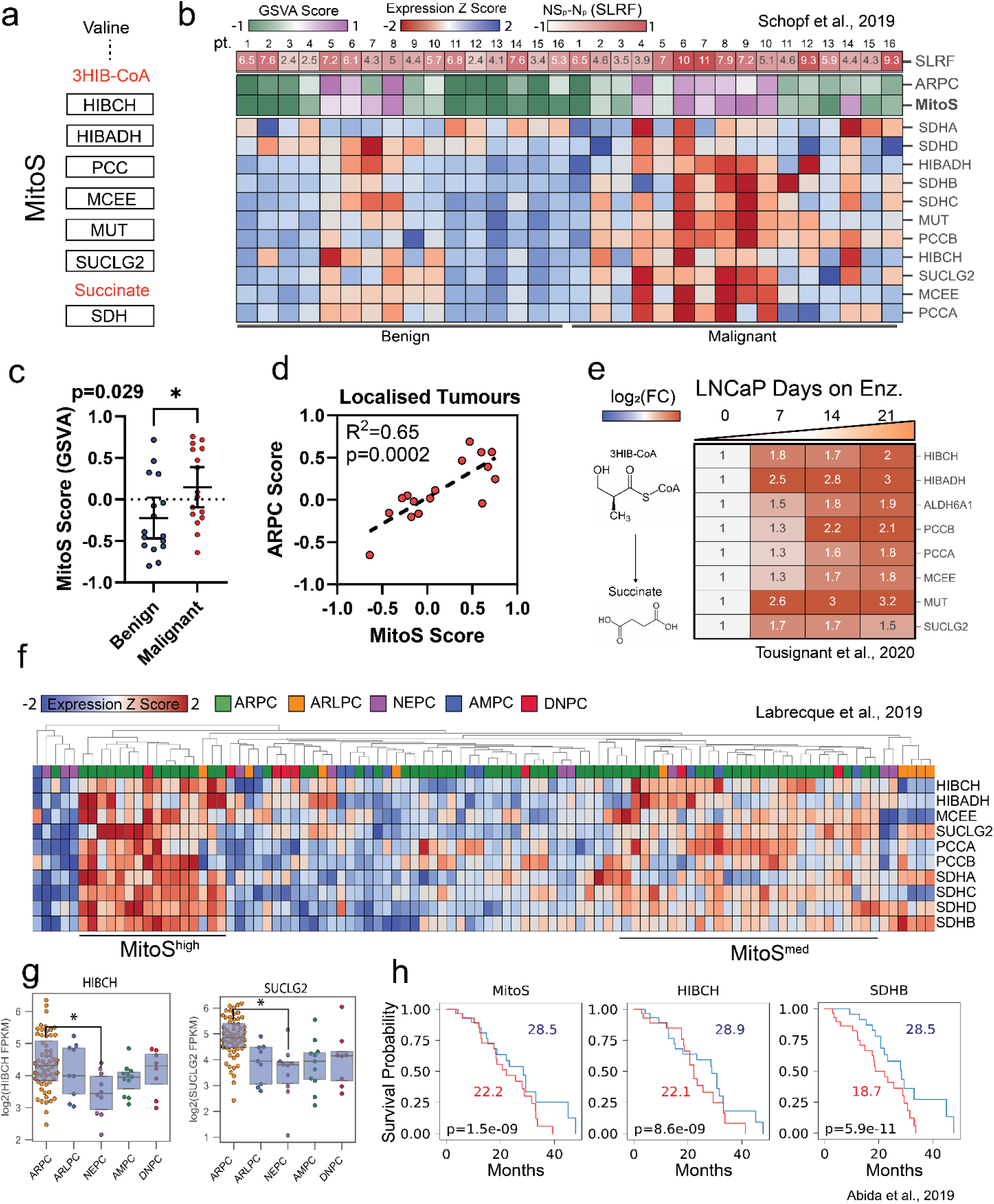
Valine Catabolism and Succinate-Linked Respiration is Enhanced Through PCa progression. (**a**) Schematic of the MitoS gene signature, generated for the purpose of this study. (**b**) Heatmap showing MitoS expression z-scores in benign vs. malignant patient primary prostate tissue with included signature scores of MitoS and AR activity (ARPC) as well as functional succinate-oxidation. (**c**) MitoS scoring by gene set variation analysis (GSVA) of benign vs. malignant prostate tissues. (**d**) Correlation analysis of MitoS and ARPC score in malignant PCa tissue. (**e**) Gene expression heatmap of enzymes responsible for 3HIB-CoA to succinate conversion in LNCaP cells treated with 7, 14 and 21 days of 10 µM enzalutamide. (**f**) Clustered heatmap of MitoS genes in metastatic castrate-resistant PCa (mCRPC) RNAseq data stratified by disease molecular subtype: AR-high PCa (ARPC), AR-low PCa (ARLPC), neuroendocrine PCa (NEPC), double-positive tumours (AMPC) and double-negative tumours (DNPC). (**g**) Gene expression of HIBCH and SUCLG2 across mCRPC subtypes. (**h**) Kaplan-Meier curves displaying the months overall survival of mCRPC patients classified into high (red) or low (blue) tumour expression of MitoS score, HIBCH and SDHB. Raw datasets were obtained from Schopf et al., 2020 (a-d), Tousignant et al., 2020 (f), Labrecque et al., 2019 (f-g) and Abida et al., 2019 (h). Significance determined by One-Way ANOVA, *p<0.05, or Kaplan Meier survival analysis.

### Inhibition of valine catabolism via HIBCH suppresses prostate cancer cell metabolism via succinate generation

To investigate the effects of inhibiting valine catabolism on PCa proliferation and succinate-linked metabolism (***Fig. 4a***), cell growth was continuously measured following an optimised siRNA-based HIBCH knockdown protocol (***Supplementary Fig. 2***). LNCaP and PC3 cell confluency (***Fig. 4b***) and cellular morphology (***Supplementary Fig. 3***) were both significantly impaired at 96 hours post-transfection. Cell proliferation was not reduced in benign BPH-1 cells, suggesting selective sensitivity in malignant prostate cells. Expression of the genes responsible for encoding the propionyl-CoA to succinyl-CoA enzymatic pathway (*PCCA, PCCB, MCEE, MUT*) were also measured by qRT-PCR at multiple time points following HIBCH knockdown. Expression of these genes was reduced at 48 hours with more substantial reductions 96 hours post-transfection. This included a significant reduction (∼50%) in *MCEE* gene expression (p=0.0004) and a non-significant reduction (∼20%) in PCCA and PCCB expression compared to siCTR (***Fig. 4c***). HIBCH protein was also measured at 72 hours by Western blot analysis post-siRNA transfection to validate knockdown (KD) which confirmed reduced expression in the BPH-1 (69% KD) LNCaP (66% KD) and PC3 (76% KD) prostate cell lines (***Supplementary Fig. 4***).

To investigate the long-term effects of HIBCH suppression, doxycycline-inducible shRNA targeting HIBCH (shHIBCH), or a non-targeting control (shNT) were generated in LNCaP cells. This model showed similar transcriptional perturbations in the genes responsible for SLRF as the siRNA model with gene expression of *SDHA* (succinate oxidation) reduced ∼50% and *SUCLG2* (succinate generation) reduced ∼40% after 72 hours dox-induction (***Fig. 4d***). We also performed 96 hours of HIBCH suppression in LNCaP cells by siRNA, which resulted in less substantial but statistically significant reductions to the expression of both the catalytic (*SDHA*: p=0.010, *SDHB*: p=0.004) and membrane bound (*SDHC*: p=0.032, *SDHD*: p=0.003) subunits of SDH (***Supplementary Fig. 5***), suggesting a sustained reduction in SLRF over time with the loss of valine catabolism. Furthermore, a statistically significant reduction to mitochondrial solute transporter *SLC25A10* expression (p=0.008) was detected, suggesting decreased shuttling of succinate and malate in response to SDH inhibition (***Supplementary Fig. 5***). To functionally confirm these results, a metabolomic analysis was conducted in LNCaP cells as a proof-of-principle experiment to demonstrate the effect of long-term (144h) shRNA mediated-inhibition of HIBCH. Substantial reductions to succinate (-39%), succinyl-CoA (-32%), fumarate (-46%), malate (-21%), cis-aconitate (-40%) and citrate (-78%) were observed within the shHIBCH samples (***Fig. 4e***). Inversely, minimal fluctuations to acetyl-CoA (-6.5%), isocitrate (-9.4%), and alpha-ketoglutarate (+5.4%) were detected. Together these results highlight the role of HIBCH in the maintenance of succinyl-CoA and succinate as well as the preservation of SDH function within the TCA cycle.

The leucine-specific enzyme methylcrotonyl-CoA carboxylase subunit 2 (MCCC2) has been previously highlighted as a therapeutic target in PCa, and was thus investigated to ensure it was not enhanced as a result of aberrant HIBCH levels^31^. Following 72 and 96 hours of HIBCH knockdown in LNCaP cells, *MCCC2* gene expression was significantly reduced (50%), suggesting a delayed reduction to leucine catabolic activity in response to suppressed valine catabolism (***Fig. 4g***). Similarly, inhibition of MCCC2 resulted in significant reductions to *HIBCH* gene expression by 48 hours post-knockdown (***Fig. 4h***). These findings provide preliminary evidence that valine and leucine catabolism are indirectly linked, and that targeting either pathway results in a delayed suppression of the other. An analysis of central metabolic gene expression further showed a statistically significant reduction in the expression of *ACCα* (p=0.0031) and GOT2 (p=0.0298), and a non-significant reduction in *CPT1α* (p=0.082) (***Fig. 4i***) Inversely, a significant increase in the expression of *ACAT1* (p=0.0005) was observed, suggesting that compensation of isoleucine catabolism may be invoked in response to reduced valine catabolic activity. Finally, non-significant increases in the expression of *HMGCR* (p=0.1862) and *HMGCS* (p=0.0897) were noted, indicating that cholesterol synthesis may be enhanced following HIBCH knockdown, which would benefit from future investigation (***Fig. 4i***).

**Fig. 4.**
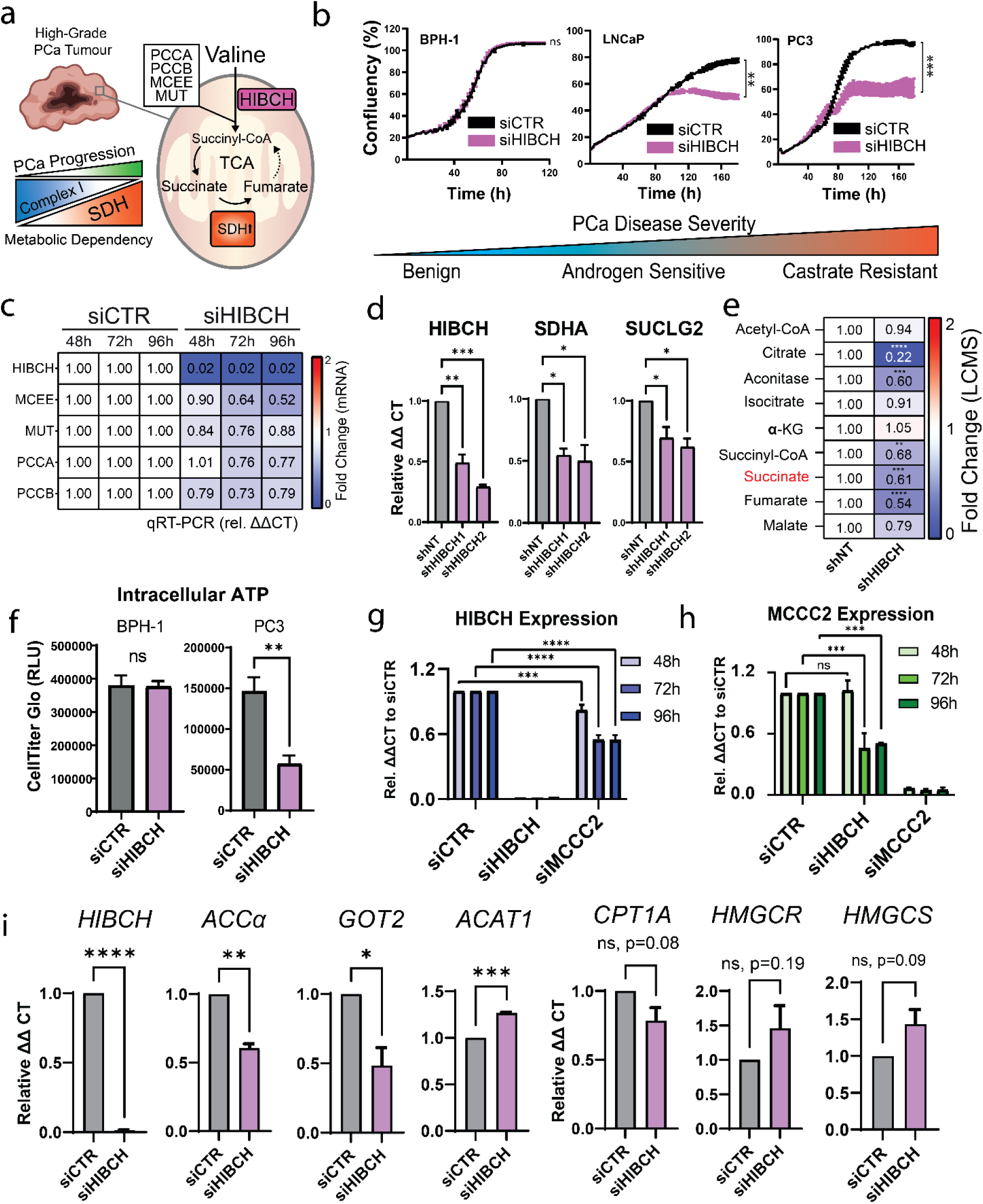
Targeting Valine Catabolism to Inhibit Proliferation and Metabolic Plasticity in PCa. (**a**) Schematic describing proposed hypothesis that mitochondrial energy remodelling (enhanced succinate oxidation by SDH) is facilitated by enhanced valine catabolism and 3-hydroxyisobutyryl-CoA Hydrolase (HIBCH) activity. (**b**) Cell confluency time course assay in LNCaP, PC3 and BPH-1 cells, following siRNA transfection of either siCTR or siHIBCH. (**c**) mRNA expression of genes encoding the succinyl-CoA generating enzymes within LNCaP cells following 24, 72 and 96 hours of siCTR or siHIBCH transfection. (**d**) mRNA expression of HIBCH, SDHA and SUCLG2 following 96 hours of induction of shCTR or shHIBCH in LNCaP cells. (**e**) Metabolomic quantification (LCMS) of tricarboxylic-acid cycle intermediates following 144 hours of induction of shCTR or shHIBCH in LNCaP cells. (**f**) ATP content of PC3 and BPH-1 cells following 96 hours of siCTR or siHIBCH transfection measured by CellTitre Glo assay. (**g**) HIBCH and (**h**) MCCC2 mRNA expression following 48, 72 and 96 hours of siCTR, siHIBCH or siMCCC2 transfection. (**i**) mRNA expression of ACCa, ACAT1, GOT2, CPT-1A, HMGCR and HMGCS by qRT-PCR following 96 hours of siCTR or siHIBCH transfection. Significance was determined by One-Way ANOVA comparing each observation to the vehicle control (siCTR or shCTR). ns – not significant, *p<0.05, **p<0.01, ***p<0.001, ****p<0.0001.

### HIBCH Knockdown inhibits mitochondrial respiration and glycolysis in PCa

To functionally measure the metabolic response of PCa cells to HIBCH knockdown, real-time metabolic flux analysis was performed with the Seahorse XFe96 Analyzer. From our previous analysis of LNCaP PCa cells, it was uncovered that valine catabolism is enhanced in response to enzalutamide. We therefore sought to compare the glycolytic and mitochondrial response of enzalutamide-sensitive (LNCaP) and enzalutamide-resistant (MR49F) cells in response to only 48 hours of HIBCH knockdown. A timeframe of 48 hours was employed in these experiments to identify acute changes in cellular respiration. LNCaP cells were exposed to either vehicle (EtOH) or 10 µM enzalutamide (ENZ) for 48 hours while MR49F cells were maintained in continuous ENZ to prevent therapeutic re-sensitization. Following treatment and siRNA transfection, glycolytic lactate export was calculated as a function of extracellular acidification rate (ECAR) while oxidative respiration was measured as a readout of oxygen consumption rate (OCR). These analyses revealed reductions to basal OCR and ECAR in both cell lines following HIBCH knockdown, however this response was muted in LNCaP cells not exposed to ENZ (***Fig. 5a***). Calculation of ATP production rate also demonstrated significant reductions to both mitochondrial and glycolytic ATP generation in these cells (***Fig. 5c***). We also noted changes to oxidative respiration were more prominent in MR49F cells, supporting the concept that enzalutamide resistance may lead to a heightened energetic dependency on HIBCH. Live cell fluorescent microscopy of MR49F mitochondria also revealed dramatic changes to mitochondrial morphology, including increased fragmentation and decreased complexity of the mitochondrial tubular network (***Fig. 5b***).

To investigate the global transcriptomic consequences (e.g. compensation) induced by HIBCH knockdown, RNA sequencing was performed on mRNA isolated from LNCaP PCa cells following 96 hours of siHIBCH transfection. Differential expression analysis was then performed on the processed dataset which revealed 497 differentially expressed genes (269 increased, 228 decreased) in response to HIBCH knockdown. Within the top 10 differentially expressed genes included a decrease of *HIBCH, UGT2B11, UGT2B28* and *SIPA1L2*, as well as the increase of *OR51E1, PLA2G2A*, *LRRN1*, *SYT4*, and *HIPK3* (***Table 1***, ***Fig. 5f***). To investigate changes to differential metabolic pathway enrichment, gene set variation scoring (GSVA) was performed. This revealed notable decrease of BCAA degradation, cysteine and methionine metabolism and expression of the ABC transporters, while nitrogen metabolism, folate biosynthesis and unsaturated fatty acid synthesis were increased (***Fig. 5d***). To improve confidence in the identification of dysregulated metabolic pathways, an integrated (joint) analysis was performed with MetaboAnalyst 5.0, to calculate an impact score derived from the previously acquired metabolomics (shHIBCH 144h) in combination with the RNAseq (siHIBCH 96h) (***Fig. 5e***). This analysis revealed that the top 5 most impacted pathways were ascorbate / aldarate metabolism, TCA cycle, pentose phosphate pathway, glycolysis and pyruvate metabolism. A limitation of the integrated analysis is that the data sets were derived from different gene silencing methods (siRNA vs shRNA), alternating time points (96h and 144h) and a targeted library of only 40 metabolites.

**Table 1.** Top 10 Differentially Expressed Genes (Up and Down) Following HIBCH Knockdown in LNCaP PCa Cells.

| Gene (Up) | Log(FC) | Adj. P Val | Ensembl99 ID | Gene (Down) | Log(FC) | Adj. P Val | Ensembl99 ID |
| --- | --- | --- | --- | --- | --- | --- | --- |
| OR51E1 | 12.294 | 2.09E-18 | ENSG00000180785 | HIBCH | -37.475 | 3.11E-65 | ENSG00000198130 |
| PLA2G2A | 8.63 | 9.28E-52 | ENSG00000188257 | UGT2B11 | -19.107 | 1.15E-79 | ENSG00000213759 |
| LRRN1 | 7.889 | 2.50E-36 | ENSG00000175928 | UGT2B28 | -13.866 | 1.01E-18 | ENSG00000135226 |
| LCN2 | 7.446 | 1.95E-05 | ENSG00000148346 | SIPA1L2 | -9.687 | 1.41E-63 | ENSG00000116991 |
| SYT4 | 6.268 | 2.04E-37 | ENSG00000132872 | ITGA1 | -6.779 | 4.41E-10 | ENSG00000213949 |
| CRYAB | 5.945 | 2.25E-04 | ENSG00000109846 | F5 | -6.462 | 1.23E-25 | ENSG00000198734 |
| HIPK3 | 5.246 | 1.39E-22 | ENSG00000110422 | AGR2 | -6.209 | 1.22E-12 | ENSG00000106541 |
| PXYLP1 | 5.132 | 7.83E-05 | ENSG00000155893 | FAR2P3 | -6.196 | 8.07E-04 | ENSG00000240253 |
| PRSS1 | 5.102 | 2.78E-03 | ENSG00000204983 | ZNF812P | -6.15 | 4.36E-16 | ENSG00000224689 |
| SLC7A2 | 4.873 | 1.77E-15 | ENSG00000003989 | ODAM | -5.867 | 5.88E-04 | ENSG00000109205 |

**Fig. 5.**
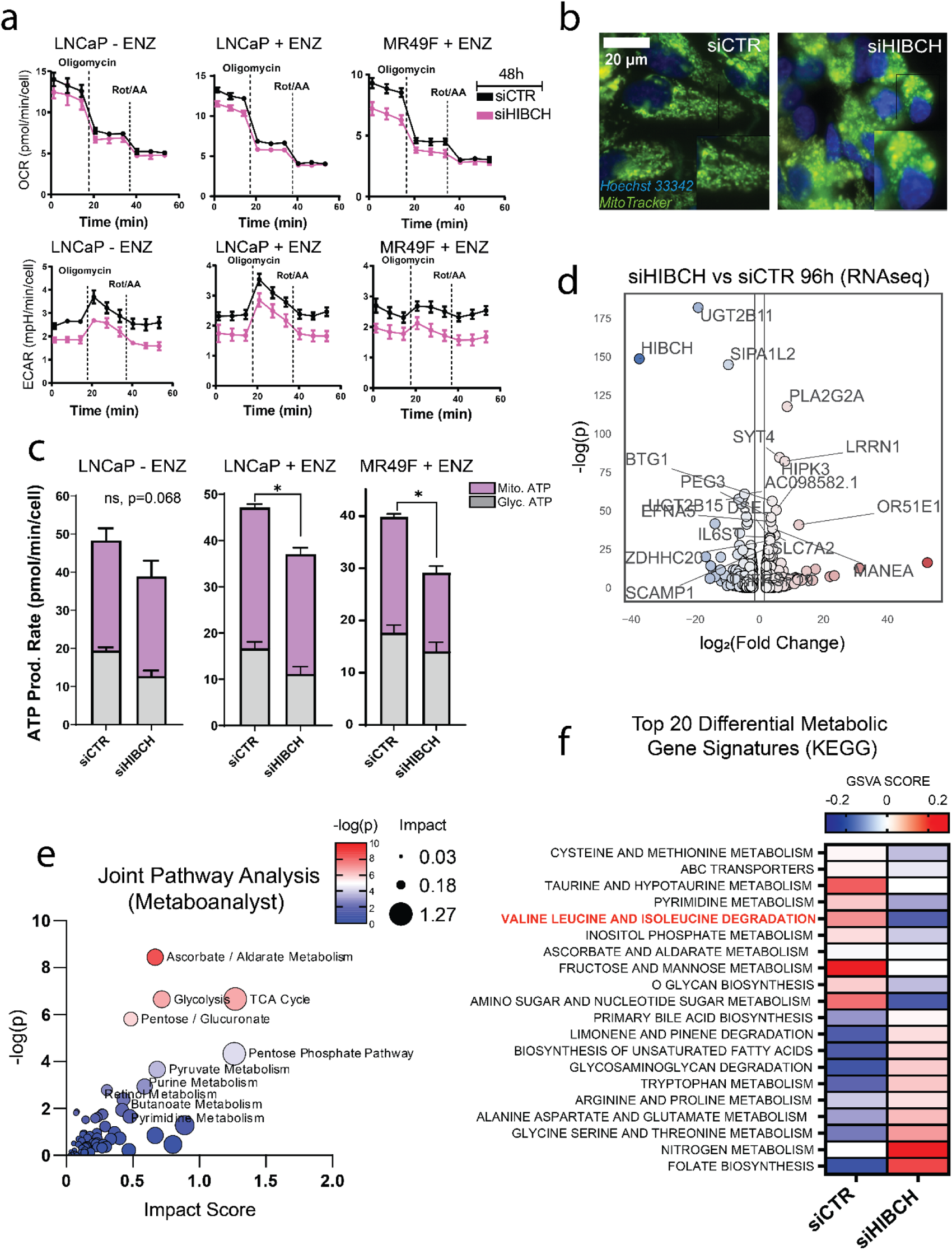
Targeting HIBCH in PCa Disrupts Oxidative Respiration, Glycolysis and Central Metabolic Pathways. (**a**) Oxygen consumption rate (OCR) and extracellular acidification rate (ECAR) following 96 hours of siCTR or siHIBCH transfection in LNCaP and MR49F cells measured by Seahorse Extracellular Flux Analysis. (**b**) Mitochondrial morphology (MitoTracker Green FM) of LNCaP cells following 72 hours of siCTR or siHIBCH transfection. (**c**) Glycolytic vs. mitochondrial ATP production rate calculated from Seahorse Extracellular Flux assay shown in Fig. 5a. (**d**)Volcano plot showing most differentially expressed genes following HIBCH knockdown. (**e**) Joint pathway analysis of both metabolomic and RNAseq data ranked by -log(p) value and impact scores. (**f**) Heatmap of the top 20 differential gene signature scores of LNCaP cells following 96 hours of siCTR or siHIBCH transfection measured by RNAseq.

## Discussion

Metabolic plasticity is an important multifaceted factor which underpins the evasive nature of cancer. We have uncovered novel insights into the role of valine catabolism in maintaining SLRF in advanced PCa. This work links the existing literature identifying the importance of BCAA degradation, lipid metabolism and oxidative phosphorylation in supporting the unique metabolic phenotype of PCa. While the BCAAs have been previously reported to be critical regulators of lipid biology within both adipose and hepatic tissues^16,32,33^, our investigation has for the first time shed light on their importance in lipid metabolism of malignant prostate cells. Our data demonstrates extracellular BCAA availability is essential for the maintenance of intracellular lipids in multiple models of advanced and castrate-resistant PCa (LNCaP, C4-2B and PC3). This finding was further supported in our BCAA catabolic inhibitory models, which demonstrated that knockdown of BCAT1 could successfully reduce lipid content in LNCaP and PC3 cells, while knockdown of BCAT2 results in an increase to lipid content in PCa cells, but not in benign BPH-1 cells. Gene expression analysis of BCAT1 and BCAT2 following their knockdown also confirmed a compensatory transcriptional relationship, highlighting a dynamic cross-regulatory mechanism. To investigate the specific roles of each BCAA, we identified an important role for valine in co-regulating fatty acid uptake in androgen responsive PCa. In response to the absence of extracellular valine, we found that LNCaP cells increase long-chain fatty acid uptake. This result is supported by other studies that link valine catabolism and fatty acid uptake through a regulatory feedback loop between HIBCH and pyruvate dehydrogenase kinase 4 (PDK4). This response was unique to PCa cells but absent in non-malignant BPH-1 cells, which instead demonstrated a unique response to leucine. Together, these investigations suggest a possible switch in metabolic dependency in PCa cells, including a modified reliance from BCAT2 to BCAT1 and from extracellular leucine to valine.

Over the past few years, enhanced succinate oxidation by SDH has been identified as an emergent hallmark of PCa metabolism^9–11^. Increased reliance on succinate in response to reduced Complex I (CI) activity is also not unique to cancer, as cell-permeable succinate has been shown to overcome CI dysfunction in models of mitochondrial disease^34^. While direct inhibition of SDH activity has been explored as a therapeutic strategy within multiple cancers due to its critical role in oxidative phosphorylation, an inherent flaw with this approach is the resultant accumulation of intracellular succinate. Increased cytoplasmic succinate is well known to stabilise hypoxia-inducible factors (HIFs), which play a pivotal role in the oncogenesis of multiple malignancies^35,36^. As a result, cells which are deficient in SDH activity are also subject to enhanced pseudohypoxia, epithelial to mesenchymal transition (EMT) and metastatic potential^37,38^. Our unique approach has aimed to resolve this problem, by instead inhibiting the upstream production of succinate via the valine catabolic axis. Our analyses of publicly available data also confirmed a transcriptomic enrichment of genes which facilitate the entry of valine into the TCA cycle (MitoS gene signature) in patient PCa tumours as well as negatively correlating to patient survival outcomes.

Inhibition of valine catabolism was achieved in this study via the suppression of HIBCH, a key enzyme facilitating the conversion of 3HIB-CoA to 3HIB. Our data demonstrates that knockdown of HIBCH results in metabolic catastrophe for PCa cells, including a reduction in intracellular succinate among other TCA metabolites, the muting of oxidative and glycolytic respiration and an induction of mitochondrial fragmentation. These findings are also supported by Shan et al (2019), who in colorectal cancer cells demonstrated that HIBCH inhibition reduces cell proliferation and TCA metabolite levels^17^. Furthermore, in response to HIBCH inhibition a time-delayed reduction to *MCCC2* gene expression was also observed, a key enzyme in the leucine catabolic pathway. MCCC2 has been previously highlighted as a therapeutic target in PCa^31^ and was thus measured to validate that leucine catabolism was not enhanced in response to HIBCH inhibition. Our findings demonstrated that not only do MCCC2 and HIBCH fail to share a compensatory relationship but are mutually dependent on the activity of the other. Our exploration into the global multi-omic changes occurring because of HIBCH inhibition also highlighted several broader metabolic consequences. This included the reduction of the genes *ACCα* and *GOT2,* as well as dysregulation of the ascorbate, cysteine/methionine, nitrogen, and folate metabolic pathways.

One knowledge gap which our study has not examined are the alternative sources of succinyl-CoA, including the catabolism of isoleucine, methionine, and threonine. However, amongst these, valine remains the most abundant and readily available amino acid in PCa patient serum (Val: 222.8 µM, Ile: 56.3 µM, Met: 16.9 µM, Thr: 102.1 µM)^39^. Our findings also highlight the significance of dissecting the specific roles for each BCAA in cellular metabolism and justify further investigations into their unique functions and regulatory mechanisms. This will be essential in further understanding the specific energetic demands which are dynamically adapted throughout the PCa disease spectrum and will assist in the generation and deployment of novel metabolically targeted strategies. More broadly, ontological pathway databases would also likely benefit from separating the leucine, isoleucine, and valine catabolic entries to prevent statistical muting introduced from signature-based analyses, which often group BCAA degradation as a singular process. Succinate has been identified as having many pro-oncogenic roles, including HIF1α stabilisation^35,36^, alteration of the tumour microenvironment^40^, promotion of metastatic potential via enhanced EMT^37^, suppression of anti-tumour immune responses^41,42^ and the activation of pro-inflammatory signalling pathways^43^. Future directions from this work would thus benefit from investigations into the targeting of these oncogenic factors via inhibition of HIBCH. In summary, our study introduces a novel therapeutic approach which exploits metabolic reliance on succinate in PCa. This multifaceted investigation reveals the intricate connections between BCAAs, lipid metabolism, and succinate-linked respiration in the context of PCa. The clinical implications of our findings highlight the potential of HIBCH inhibition as a targeted therapeutic strategy for PCa, warranting further exploration and validation.

## Methods

### Cell Culture

Benign and malignant prostate cell lines: BPH-1, LNCaP, C4-2B, PC3 and MR49F were cultured in RPMI 1640 (Gibco) supplemented with 5% foetal bovine serum (FBS). The enzalutamide-resistant cell line MR49F was continuously maintained in 10% FBS + 10µM enzalutamide (MedChemExpress) to prevent therapeutic re-sensitisation. Medium was replenished at least every 72 hours and cells were routinely screened for mycoplasma.

### Amino Acid Depletion Assays

Following 72 hours of standard culture in poly-L-ornithine coated 96-well optical plates (Cellvis, #P96-1.5H-N), culture medium was washed twice and refreshed with BCAA depleted medium (BDM). BDM was generated though supplementation of glucose and amino acids (except BCAAs) to RPMI 1640 Medium Modified without L-Glutamine, Amino acids, Glucose or Phenol Red (MyBioSource, #MBS653421). BDM was supplemented with 10% dialysed FBS to prevent the addition of amino acids from serum. Exogenous supplementation of leucine (100 µM), isoleucine (55 µM) and valine (220 µM) were then performed to restore physiologically representative BCAA serum concentrations. The formulation of BDM has been included in the Supplementary Data.

### Small Interfering RNA (siRNA) Knockdown

siRNA silencing was accomplished using forward transfection of pre-designed MISSION siRNAs from Sigma-Aldrich. A 10 nM final concentration of siBCAT1 (SASI_Hs01_00066058), siBCAT2 (SASI_Hs02_00331330), siHIBCH (SASI_Hs01_00064760) or siMCCC2 (SASI_Hs01_00039625) was transfected into cells with RNAiMAX transfection reagent (ThermoFisher, #13778150) diluted 1:25 in serum free RPMI 1640. Following 5 hours of transfection, standard serum conditions were restored by the addition of FBS (5%). A universal negative control (Sigma Aldrich, #SIC001) and Cy5-labelled negative control (Sigma Aldrich, #SIC005) with no known human homologies were included in all experiments to control for non-specific toxicity and validate successful lipofection.

### Inducible Short Hairpin RNA (shRNA) Knockdown

Glycerol *Escherichia coli* stocks containing SMARTvector inducible lentiviral shRNA (mCMV, tGFP) were obtained from Horizon Discovery. Purified plasmid DNA was generated from overnight LB cultures with the QIAprep Spin Miniprep Kit (Qiagen, #27106X4) and quantified using the NanoDrop ND-1000 spectrophotometer (ThermoFisher, #ND-1000). Viral supernatants were generated in 10 cm dishes by transfection of HEK293T cells with 0.2 µg pCMV delta R8.2 (lentiviral packaging plasmid), 1.8 µg pCMV-VSV-G (lentiviral envelope plasmid) and 2 µg purified shRNA DNA with 12 µg (3:1) of X-tremeGENE transfection reagent (Sigma Aldrich, #6366244001). Following 72 hours of transfection, LNCaP cells were transduced with fresh viral supernatant for 6 hours and the cultures expanded. Target-specific regions of shHIBCH1 and shHIBCH2 were ***GAATAATGCTGTTGGTTCT*** and ***ACTCTTAATATGATTCGGC,*** respectively.

### Quantitative Reverse Transcription PCR (qRT-PCR)

Following cell siRNA transfection or shRNA induction, total RNA was isolated using the Total RNA Purification Plus Kit (Norgen, #48300) according to the manufacturer’s protocol. RNA concentration was quantified using the NanoDrop ND-1000 (ThermoFisher, #ND-1000) and RNA purity was subsequently assessed. cDNA was generated from 1 µg of RNA using the SensiFAST cDNA synthesis kit (Bioline, #BIO65054). cDNA was then diluted (1:5) with RNAase/DNAase-free water and qRT-PCR performed using SYBR-Green Master Mix (ThermoFisher, #4309155) and primers listed in Supplementary Data. qPCR was monitored on the ViiA-7 Real-Time PCR system (Applied Biosystems). Relative mRNA expression was then calculated using the ΔΔCT method normalising gene expression to the RPL32 house-keeping gene.

### SDS-PAGE and Western Blotting

Cells were scraped and lysed with 100 µL of freshly prepared RIPA buffer. Samples were centrifuged at 14,000 RCF (x g) for 15 minutes before collection of the supernatant and removal of cellular debris. Protein concentrations were measured using the Pierce BCA Protein Assay Kit (ThermoFisher, #23225). 50 µg of protein lysate was prepared with 4X Bolt™ LDS Sample Buffer (ThermoFisher, #B0007) and 10X Bolt™ Sample Reducing Agent (ThermoFisher, #B0009) and loaded onto a Bolt™ 4-12% Bis-Tris Mini Protein Gel (ThermoFisher, #NW04127). Gel electrophoresis was run at 120 V (constant) for 60 minutes in 1X MOPS SDS running buffer (ThermoFisher) using the Bolt Mini Blot Module (ThermoFisher, #B1000). Protein transfer using nitrocellulose membrane was conducted in 1X Bolt Transfer Buffer at 10V (ThermoFisher, #BT0006) for 1 hour, then blocked in Odyssey TBS Blocking Buffer (LI-COR, #927-500) for 1 hour at room temperature. HIBCH and y-Tubulin antibodies were diluted 1:1000 in blocking buffer and incubated with the membrane overnight at 4°C. The membrane was washed again and then incubated with 1:25000 Odyssey fluorescent-dye conjugated secondary antibodies (LI-COR) for 1 hour at room temperature protected from light. The membrane was again washed and imaged using the Odyssey infrared imaging system (LI-COR) and densitometry quantified using Image Studio Lite (LI-COR).

### Intracellular Lipid Content Quantification

Following 24 hours of culture in BDM, media was removed, and the optical plate was washed twice with 200 µL PBS. Cells were fixed with 4% paraformaldehyde (PFA) (Electron Microscopy Sciences, #50-980487) at room temperature for 30 minutes, rinsed with PBS, then intracellular lipids were co-stained with 0.1 µg/mL Nile Red (Invitrogen, #N1142) and 1 µg/mL 4,6-diamidino-2-phenylindole (DAPI) (Invitrogen, #D1306) in PBS for 1 hour at 4°C. Automated fluorescent microscopy was accomplished using the InCell Analyzer 6500HS (Cytiva, #6500HS). Ten fields per well were captured using the 405, 561 and 642 nm (Phospholipids) excitation filters at 10X and 60X magnifications. Image segmentation and quantitation of cellular parameters from fluorescent 10X imaging were derived using custom pipelines in CellProfiler 4^18^. Representative images (60X) where processed with ImageJ and identical thresholding settings were applied across all images where comparisons have been made^19^.

### Live Cell C16:0-BODIPY Uptake Assay

Following 24 hours of culture in BDM, media was removed, and the optical plate was washed twice with 200 µL PBS. Cells were then treated with 5 µM C16:0 BODIPY (C16) (Invitrogen, #D3821), 1 µg/ml Hoechst 33342 (ThermoFisher, #62249) and 0.2% bovine serum albumin (BSA) (Sigma Aldrich, #A8412) in serum-free BDM for 30 minutes at 37°C to allow cellular uptake of C16. Cells were washed twice with 200 µL serum-free BDM before imaging. Images were captured using the InCell Analyzer 6500HS (Cytiva). Ten fields per well were captured with the 405 nm (Blue, DAPI) and 488 nm (Green, C16) excitation filters at 10X and 60X magnifications. Image analysis was performed as described above.

### Seahorse Extracellular Flux Assay

Quantification of glycolytic and mitochondrial ATP production was performed using the Seahorse XFe Real-Time ATP Rate Assay Kit (Agilent, #103591-100). Prior to cell adhesion, Seahorse 96XFe tissue culture plates were coated with 0.01% poly-L-ornithine solution (Sigma-Aldrich, #P4957) and incubated at 37°C. Cells were seeded at a density of 10,000 cells per well in standard phenol red-free RPMI 1640 (ThermoFisher, #11835055) with 5% FBS. Cells were subject to 48 hours of siRNA mediated HIBCH knockdown as described above. At endpoint, the media was replaced with Seahorse Base Medium (Agilent, #103334-100), supplemented with 2 mM L-glutamine (Sigma Aldrich, #1294808), 10 mM glucose (ThermoFisher, #A2494001) and 1 mM sodium pyruvate (ThermoFisher, #11360070) adjusted to pH 7.4. Oligomycin (15 µM) and a combined injection of 10 µM antimycin A + 10 µM rotenone were added to the appropriate Seahorse XFe injection ports. The Seahorse XFe96 analyser was programmed to measure both oxygen consumption rate (OCR) and extracellular consumption rate (ECAR) over the course of 60 minutes. Nine measurements were taken at 7-minute intervals with oligomycin injected between measurements 3 and 4 and rotenone + antimycin A injected between measurements 6 and 7. Extracellular flux data was analysed using Wave desktop software (Agilent) and normalised to the number of cells in each well quantified with a Hoechst nuclear stain.

### LCMS Metabolite Quantification

HIBCH knockdown was induced in shHIBCH and shNT LNCaP cells with 0.25 µg/mL doxycycline for 144 hours. Cells were rapidly washed with 5 mL isotonic wash solution (5% D-mannitol) on ice to remove residual media and exogenous metabolites without disturbing metabolite integrity as previously described^20^. Instantly, 1 mL of ice-cold extraction buffer (50% methanol + 250 nM AZT standard) was added before cells were scraped and transferred into to 15 mL conical tubes. This step was again repeated, and the two fractions were combined. Samples were freeze-thawed (-80°C) then sonicated on ice for 10 minutes. Metabolites were purified from the organic fraction of a phenol:chloroform:isoamyl (25:24:1) solution and centrifuged at 16,000 RCF for 5 minutes at 4°C. The supernatant was removed without disturbing the interface layer, frozen and sent to the Metabolomics Australia Queensland Facility for analysis. Samples were freeze-dried and resuspended in 100 µL 2% acetonitrile (ACN) before being injected (LCMS) in two dilutions to account for natural variability in metabolite levels.

### RNA Sequencing

Total cellular RNA from LNCaP cells following 96 hours of HIBCH knockdown was extracted using the Norgen RNA Purification PLUS kit (Norgen, #48400) according to the manufacturer’s instructions. RNA quality and quantity were determined on an Agilent 2100 Bioanalyzer (Agilent Technologies, Santa Clara, USA) and Qubit® 2.0 Fluorometer (ThermoFisher). Library preparation and sequencing was performed at the QUT Central Analytical Research Facility (CARF) using the Illumina TruSeq Stranded mRNA Sample Prep Kit (strand-specific, polyA enriched, Illumina, San Diego, USA) with an input of 1 µg total RNA (RIN>9), followed by paired-end sequencing on the Illumina NovaSeq6000. Raw reads were trimmed using TrimGalore, followed by alignment to the human genome (GRCh38 / hg38) using the STAR2 aligner and read quantification with RSEM4^21,22^. Differential expression (DE) between siCTR and siHIBCH samples was calculated after sample normalization using edgeR (no replicates: Fisher Exact Test; replicates: General Linear Model) and is defined by an absolute fold change of >=1.5 and an FDR corrected p-value <=0.05^23^. Data quality control included running FastQC before and after trimming, checking RNAseq metrics with the PICARD tool kit, and mapping reads against microbial genomes using Kraken^24,25^.

### Statistics and Bioinformatic Analyses

Statistical analyses were performed either with GraphPad Prism 10 (GraphPad Software) or the SciPy python package. Statistical tests are described in the figure legends^26^. Hierarchically clustered heatmaps were produced using the seaborn python package. Kaplan Myer survival plots were generated using the lifelines python package. Gene signature scoring was accomplished using Gene Set Variation Analysis (GSVA) package in R, incorporating the respective transcript per million (TPM) values of genes listed in the Supplementary Data^27^. Joint pathway analysis was achieved in MetaboAnalyst 5.0 using differentially expressed fold change values (FC>=1.5, p=<0.05) derived from the RNAseq and metabolomic experiments listed above^28^.

