## Supplementary Figures for "Valine Catabolism Drives Bioenergetic and Lipogenic Fuel Plasticity in Prostate Cancer"

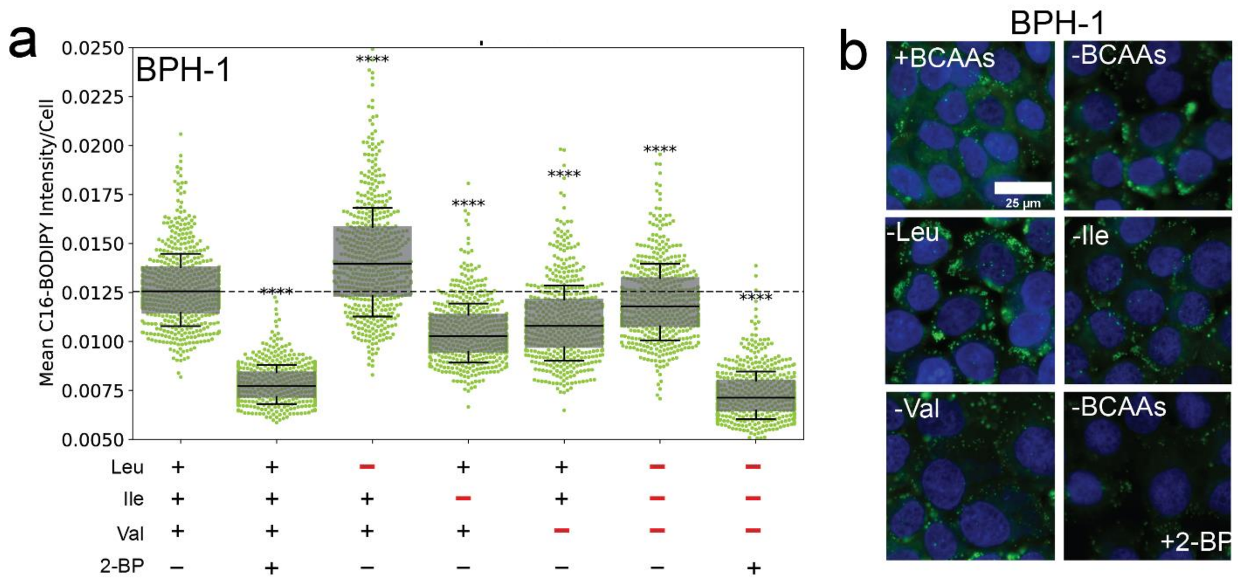


***Supplementary Fig. 1: Leucine Deprivation Triggers C16:0 Fatty Acid Uptake in Benign BPH-1 Cells**. Quantitative single cell imaging (qSCI) analysis **(a)** and representative live-cell fluorescent images **(b)** of C16-BODIPY uptake following 24h of exogenous BCAA deprivation and/or co-treatment with 2-Bromopalmitate 2-BP in BPH-1 cells.*

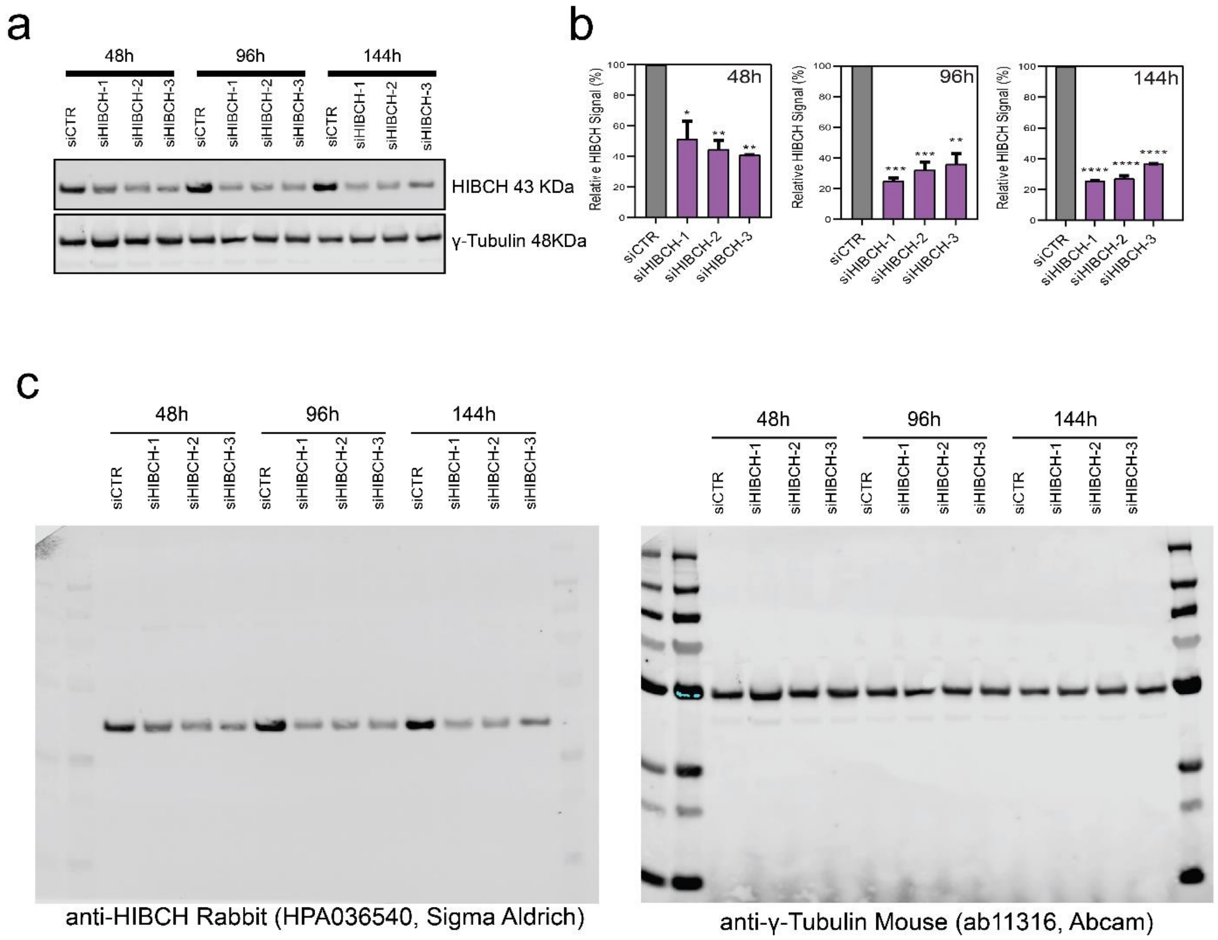


***Supplementary Fig. 2: Optimisation and Confirmation of HIBCH knockdown by siRNA Transfection*** *Comparison of three HIBCH siRNAs was performed in LNCaP PCa cells to determine the most effective and stable siRNA sequence. siHIBCH-1 (MISSION siRNA #SASI_Hs01_00064760) resulted in the highest knockdown efficiency at 144h in LNCaP cells quantified by western blot (**a**-**b**) and was thus selected for use within the present study. Full western blots with antibody catalogue numbers are included above (**c**).*

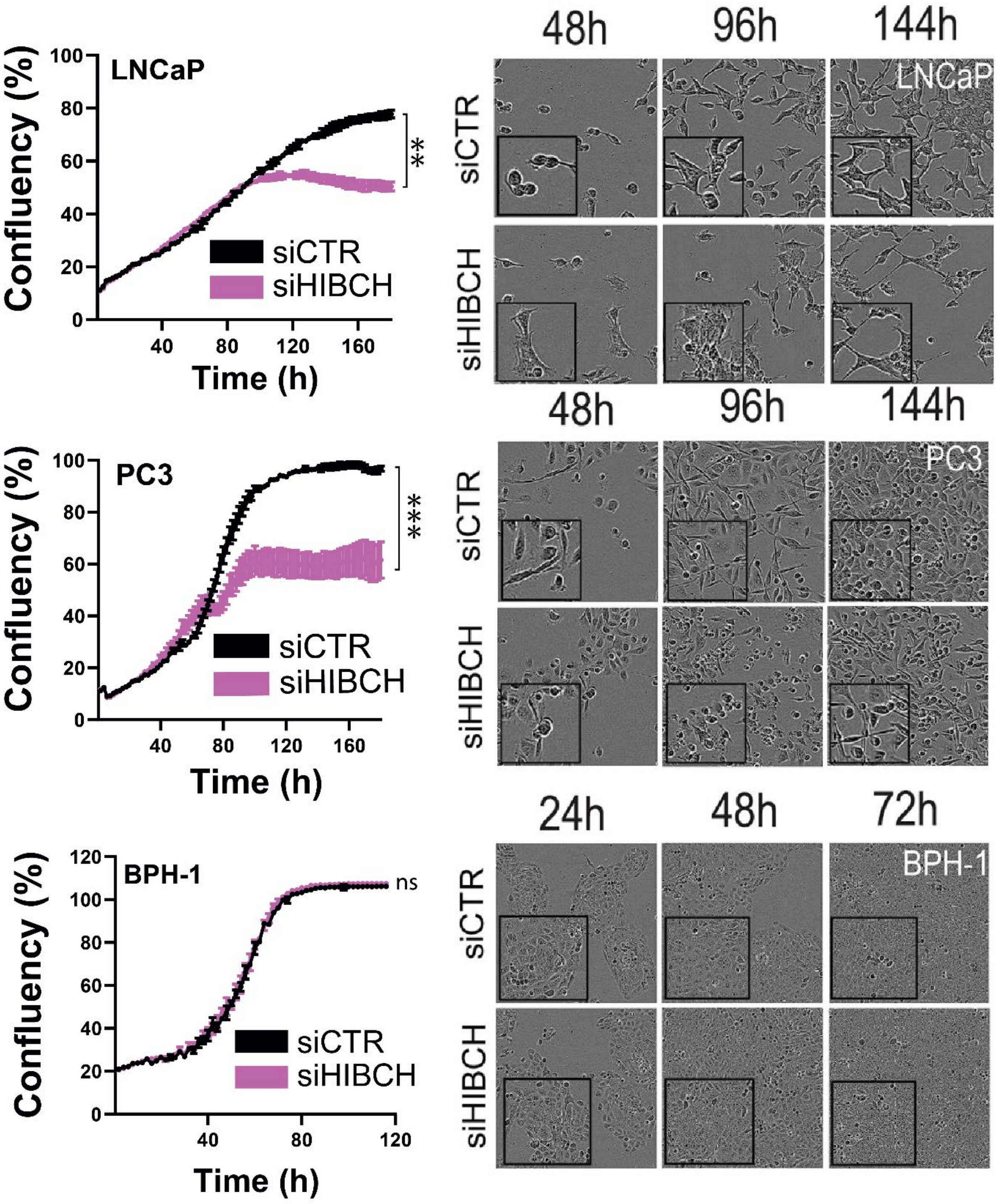


***Supplementary Fig. 3: PCa cell morphology is altered following HIBCH knockdown.***
*Cell confluency time course assay and respective images of LNCaP, PC3 and BPH-1 cells, following siRNA transfection of either siCTR or siHIBCH taken with the IncuCyte S3 Live Cell Analyzer.*

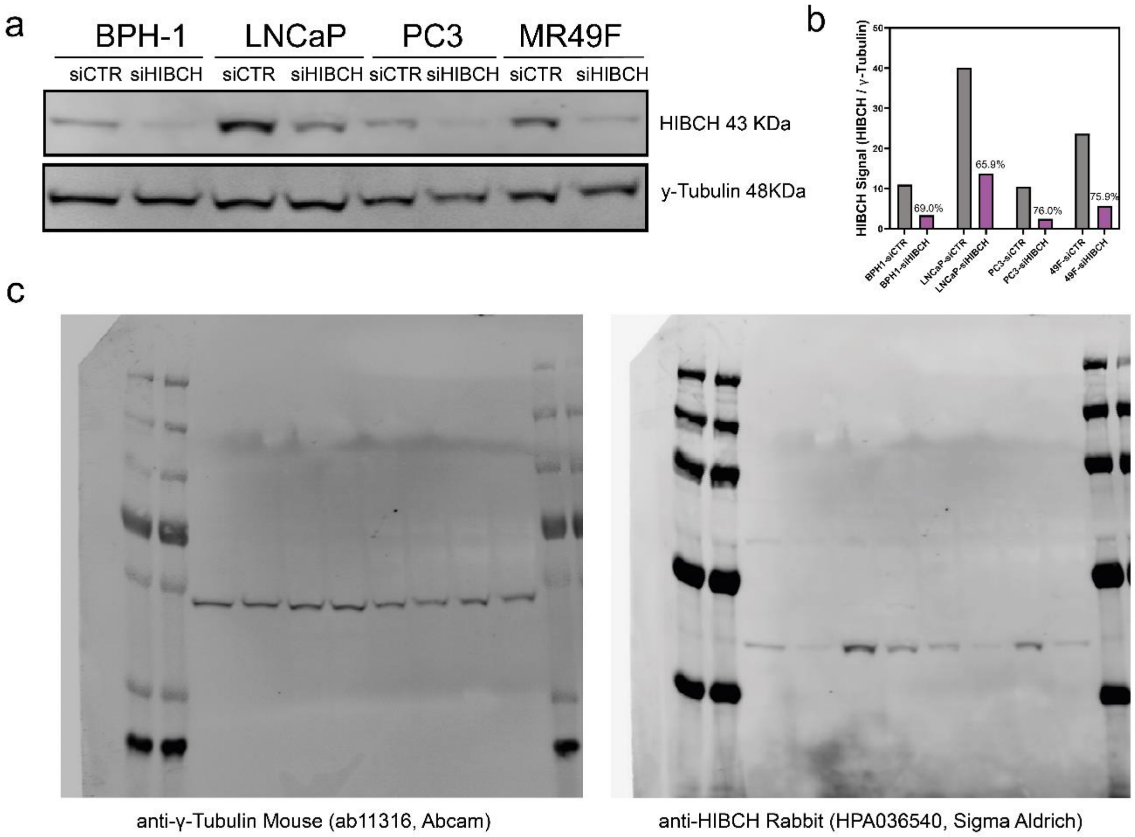


***Supplementary Fig. 4: Validation of HIBCH knockdown in LNCaP, PC3, BPH-1 and MR49F cells.***
*(**a**) Western Blot and (**b**) densitometric analysis of HIBCH and y-Tubulin protein in PCa cell lines following 96 hours siRNA (siCTR or siHIBCH) transfection. (**c**) Full western blots with antibody catalogue numbers.*

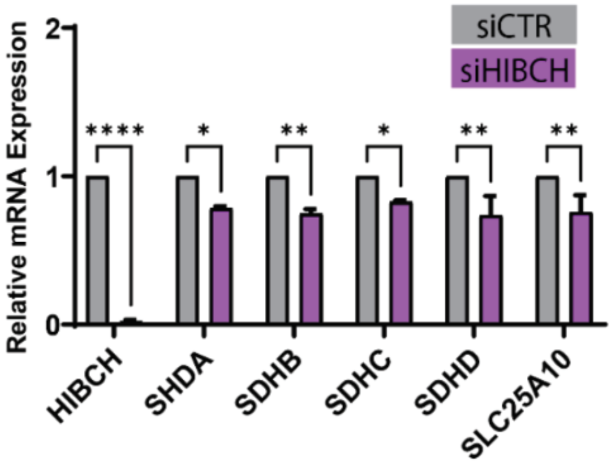


***Supplementary Fig. 5: siRNA mediated suppression of HIBCH reduces succinate-related genes.*** *Gene expression of SDH subunits (A-D) and succinate/malate transporter SLC25A10 at 96 hours post-transfection measured by qRT-PCR analysis.*
